# The impact of sex on gene expression in the brain of schizophrenic patients: a systematic review and meta-analysis of transcriptomic studies

**DOI:** 10.1101/2023.02.13.528356

**Authors:** Hector Carceller, Marta R. Hidalgo, Gonzalo Anton-Bernat, María José Escartí, Juan Nacher, Maria de la Iglesia-Vayá, Francisco García-García

## Abstract

**Background:** Schizophrenia is a severe neuropsychiatric disorder characterized by altered perception, mood, and behavior that profoundly impacts patients and society despite its relatively low prevalence. Previous studies have suggested that the dopamine D2 receptor gene and genes involved in glutamatergic neurotransmission, synaptic plasticity, and immune function as genetic risk factors. Sex-based differences also exist in schizophrenia epidemiology, symptomatology and outcomes; however, we lack a transcriptomic profile that considers sex and differentiates specific cerebral regions.

**Methods:** We performed a systematic review on bulk RNA-sequencing studies of post-mortem brain samples. Then, we fulfilled differential expression analysis on each study and summarized their results with regions-specific meta-analyses (prefrontal cortex and hippocampus) and a global all-studies meta-analysis. Finally, we used the consensus transcriptomic profiles to functionally characterize the impact of schizophrenia in males and females by protein-protein interaction networks, enriched biological processes and dysregulated transcription factors.

**Results:** We discovered the sex-based dysregulation of 265 genes in the prefrontal, 1.414 genes in the hippocampus and 66 genes in the all-studies meta-analyses. The functional characterization of these gene sets unveiled increased processes related to immune response functions in the prefrontal cortex in male and the hippocampus in female schizophrenia patients and the overexpression of genes related to neurotransmission and synapses in the prefrontal cortex of female schizophrenia patients. Considering a meta-analysis of all brain regions available, we encountered the relative overexpression of genes related to synaptic plasticity and transmission in female and the overexpression of genes involved in organizing genetic information and protein folding in male schizophrenia patients. The protein-protein interaction networks and transcription factors activity analyses supported these sex-based profiles.

**Conclusions:** Our results report multiple sex-based transcriptomic alterations in specific brain regions of schizophrenia patients, which provides new insight into the role of sex in schizophrenia. Moreover, we unveil a partial overlapping of inflammatory processes in the prefrontal cortex of males and the hippocampus of females.

**Plain language summary:** Schizophrenia is a severe neuropsychiatric disorder characterized by altered perception, mood, and behavior that profoundly impacts patients and society. Previous studies have suggested dopamine and glutamate neurotransmission genes, as well as immune function alteration as genetic risk factors. Schizophrenia epidemiology, symptomatology and outcomes are different for women and men, but the biological reason is not understood. Therefore, we reviewed all RNA-sequencing studies of post-mortem brain samples of women and men affected by schizophrenia available. Then, we compared the gene expression on each study for males and females and integrated the results of studies on different regions meta-analyses: prefrontal cortex, hippocampus and all-studies. Finally, we functionally characterize the impact of schizophrenia in males and females by protein-protein interaction networks, enriched biological processes and dysregulated transcription factors. We discovered the sex-based dysregulation of 265 genes in the prefrontal cortex, 1.414 genes in the hippocampus and 66 genes in the all-studies meta-analyses. The functional characterization of these genes unveiled increased immune response functions in the prefrontal cortex in men and the hippocampus in women schizophrenia patients, as well as increased neurotransmission and synapses in the prefrontal cortex of female schizophrenia patients. The protein-protein interaction networks and transcription factors activity analyses supported these sex-based profiles. Our results report multiple transcriptomic alterations in specific brain regions of schizophrenia patients, which provides new insight into the role of sex in schizophrenia. Moreover, we unveil a partial overlapping of inflammatory processes in the prefrontal cortex of males and the hippocampus of females.

**Highlights:**

- The expression of 265 genes is altered in the prefrontal cortex of schizophrenic patients, being overexpressed in females those related to synaptic transmission.
- In the prefrontal cortex of males, overexpressed genes and overactivated transcription factors are linked to immune response and inflammation.
- Conversely, genes and transcription factors more activated in the hippocampus of females are related to immune response, whereas those genes more expressed in males are linked to protein processing.
- The global meta-analysis unveils groups of long non-coding genes and pseudogenes differentially expressed in males and females.
- The effects of schizophrenia are closely related in the prefrontal cortex of males and the hippocampus of females.

## Introduction

Coined in 1908 and affecting approximately 1% of the global population, the term “schizophrenia” describes a psychiatric disorder resulting from an intricate interaction of genetic and environmental factors that alters brain development [1,2]. The onset of schizophrenia generally occurs around late adolescence, with symptoms classically divided into positive, negative, and cognitive categories and outcomes ranging from total recovery to severe chronic impairment [2,3].

Multiple studies of the molecular pathogenesis of schizophrenia have encountered alterations in three neurotransmission systems (dopaminergic [4], glutamatergic [5], and GABAergic [6]) in the post-mortem brain; additionally, genome-wide association studies (GWAS) support these findings [7].

Multiple sex-based differences have been described in schizophrenia epidemiology. While females suffer from an older average age of onset [8], males suffer from a slightly higher incidence [9]; furthermore, studies have described a higher frequency of positive, affective symptoms in females with more severe negative symptoms in males [10]. Unfortunately, the molecular mechanisms underlying noted sex-based differences remain relatively unexplored, which has hampered the identification of sex-specific biomarkers and therapeutic interventions.

In-silico approaches such as meta-analyses of transcriptomic data have the potential to unveil novel associations by integrating multiple data sources; however, previous studies addressing sex-based differences in gene expression in schizophrenia patients (mainly focusing on the prefrontal cortex or PFC) have yet to reveal the significant impact of biological sex on gene expression [11–13].

We carried out a comprehensive systematic review of available studies in public repositories to evaluate sex-based differences in transcriptomic data from male and female schizophrenic patients. We performed three meta-analyses of gene expression in the PFC, the hippocampus, and the whole brain and encountered region- and sex-specific gene expression profiles and a partial overlap of schizophrenia-associated alterations between the hippocampus in female and the PFC in male schizophrenia patients. Moreover, functional enrichment of differentially-expressed genes (DEG) highlighted this overlap, both using gene set expression analysis (GSEA) and transcription factor activity analysis. Our results suggest significant sex-based differences in gene expression profiles in male and female schizophrenia patients, including a partial consensus in gene expression profiles found in the female hippocampus and male PFC.

## Material and Methods

### Systematic Review

Following PRISMA statement guidelines [14], a systematic review was performed by searching for the keyword “schizophrenia” in multiple databases (Gene Expression Omnibus (GEO), Array Express) during February of 2022 (revised period: 2002-2022). We also conducted a search using Google for the keywords “schizophrenia”, “RNAseq” and “human”. Two researchers (HC and FGG) independently screened titles and abstracts of all articles retrieved. Results were filtered by i) organism: “*Homo sapiens*,” ii) study type: “expression profiling by high throughput sequencing,” and iii) sample count: at least twelve samples. The exclusion criteria applied were: i) experimental design other than patients vs. controls, ii) the absence of information on patient sex, iii) experimental samples other than post-mortem brain samples, iv) the absence of female or male patient samples, and v) experimental data excluding cellular populations. To apply these criteria, HC screened full-text articles to obtain the required information. If this information was unclear, we contacted authors to provide further details. Therefore, gene expression data of five RNA-sequencing (RNA-seq) datasets were retrieved from GEO database: GSE174407, GSE138082, GSE80655, GSE42546, and GSE78936. Additionally, the raw sequence reads files of the SRP102186 study were downloaded from the Sequence Read Archive (SRA) database, whose expression dataset is unavailable. We assessed risk of bias in the included studies by the reanalysis of raw data following a common workflow detailed in the next section.

### Analysis

The workflow for each of the selected studies was: i) data download and normalization, ii) exploratory analysis of data, and iii) analysis of DEG. Then, DEG results were integrated into three meta-analyses based on the region of study -the 1) PFC, 2) hippocampus, and 3) whole brain - and functional enrichment of results was performed using protein-protein interactions (PPI), gene set expression analysis (GSEA), and transcription factor activity analysis. Studies were grouped for meta-analysis based on the brain region, allowing the exploration of common sex-based differences throughout multiple brain regions and those specifically present in the PFC and hippocampus. Bioinformatics analysis used R and all code used is available in Zenodo repository (http://doi.org/10.5281/zenodo.7778277).

### Data Download and Processing

Raw expression data of selected studies were downloaded and standardized as follows: conversion and updating (if necessary) gene names to Ensembl gene ID nomenclature, calculation of the median expression in the case of multiple values for a gene, and log2 normalization of expression data on those studies with raw data. In those studies that included multiple regions (GSE138082 and GSE80655), expression data was split to achieve one data frame per region. For the SRP102186 study, raw FASTQ files were downloaded, quality control was performed with FastQC [15] and aligned against the reference human genome GRCh38.p13 using Hisat2 [16]. The results were sorted with SAMtools [17], and raw counts were obtained with HTSeq [18]. Regarding metadata, labels used for the sex of patients were homogenized, and individuals were grouped into ‘Control’ or ‘Schizophrenia.’ After data normalization, batch effects or anomalous behavior of data were analyzed using clustering and principal component analysis. We also collected all the available data about patients characteristics: sex, age, smoking status, cause of death and drug treatments.

### Differential Gene Expression and Meta-analyses

DGE analyses for each selected study were performed using the voom function of the R package limma [18,19], adding the control factor ‘age’ to the model. Specifically, the schizophrenia sex-differential impact was analyzed through the comparison ((Schizophrenia.Female -Control.Female) -(Schizophrenia.Male -Control.Male)). The logarithm of the fold change (LFC) was used to measure the statistical effect in the DEG analysis. Thus, genes with positive LFC were increased in females, while genes with a negative LFC were increased in male schizophrenia patients. After calculating the differential expression statistics, p-values were adjusted using the Benjamini & Hochberg (BH) method. The DEG results of each study were then integrated using the metafor R package, as previously described [20]. The sex-based comparison and brain region under study (PFC, hippocampus, and whole brain) were then meta-analyzed using a random-effects model for each gene using LFC as an expression measure and standard error as a variance measure [21]. The meta-analyses provide a p-value adjusted by the BH method. The significance cutoff was set to p-value < 0.05 and absolute LFC > 0.5. Finally, DEGs previously related to schizophrenia in the Open Targets database were explored [22].

### Functional Enrichment Analysis

PPI networks were analyzed using the STRING web tool for each DEGs subset [23]. The total number of edges was examined, and PPI enrichment was assessed using the following parameters: 1) for active interaction sources, the options ‘Text Mining’ and ‘Databases’ were excluded, and 2) the minimum required interaction score was set to 0.7 (high confidence). The rest of the parameters were set as default. For enriched PPI networks, we manually colored the elements of connected components.

Then, to detect sex-based functions affected by schizophrenia, a GSEA of the three meta-analyses was performed using the logistic regression model implemented in the mdgsa R package [24]. The functional annotation of biological processes (BP) was obtained from the Gene Ontology (GO) database [24,25]. Due to their hierarchical structure, gene annotations were propagated with GO terms. Therefore, excessively specific or generic annotations were filtered out (blocks smaller than 10 or larger than 500 genes). Finally, function p-values were adjusted with the Benjamini & Yekutieli (BY) method, and the logarithm of the odds ratio (LOR) was used to measure the statistical effect. The significance cutoff was set to p-value < 0.05.

Finally, transcription factor activity was estimated based on the consensus LFC of each gene meta-analysis and its alteration by schizophrenia. The DoRothEA package [26] was used to select 271 regulons with a high confidence level (A, B, and C) of the *Homo sapiens* network. Then, protein activity was inferred using VIPER [26], obtaining a normalized enrichment score to measure transcription factor activity, and the p-value was adjusted using the BH method.

### Metafun-SCZ web tool

All data and results generated in the different steps of the meta-analysis are freely available on the Metafun-SCZ platform (http://bioinfo.cipf.es/metafun-SCZ) to any user, allowing the confirmation of obtained results and the exploration of other results of interest. This easy-to-use resource is divided into four sections: i) the summary of analysis results in each phase, followed by detailed results for the ii) exploratory analysis, iii) meta-analysis, and iv) functional profiling for each study. The user can interact with the web tool through graphics and tables and explore information associated with specific genes or biological functions.

## Results

### Data Acquisition and Preliminary Analyses

Our systematic review screened 96 publications and found eight includible studies (**Figure 1**). Finally, we discarded [27] and [28] because they did not include all cellular populations from the original sample. From selected studies [29–34], two contained expression data from three different regions. Therefore, we split these data frames by region to obtain ten datasets: four PFC, four hippocampus, one amygdala, and one nucleus accumbens (**Supplementary Table 1**).

**Figure 1.**
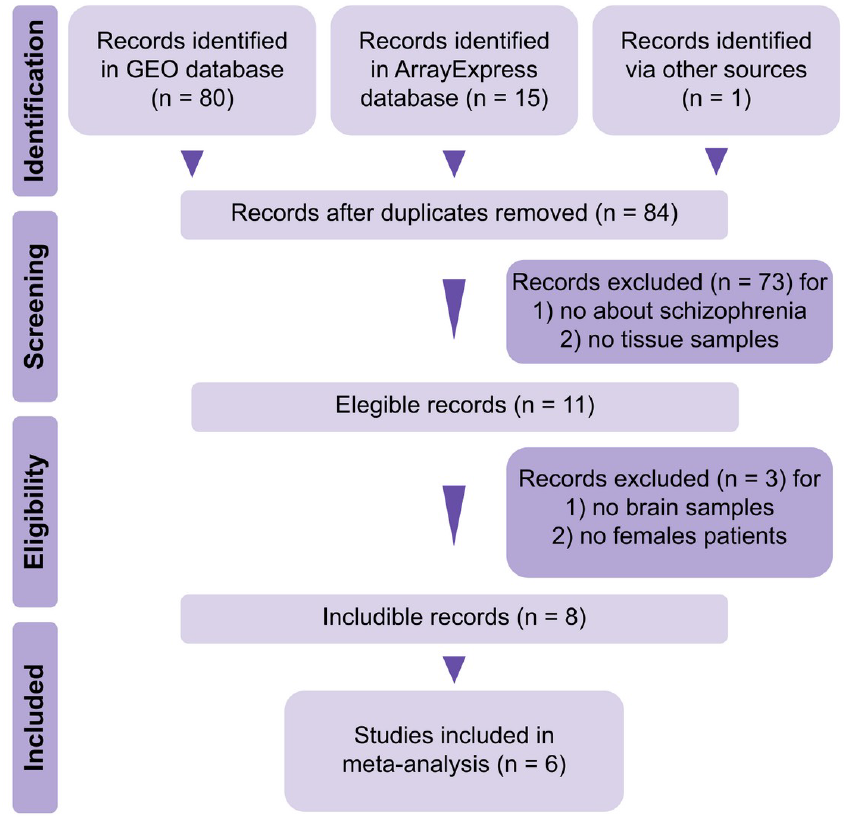
Prisma diagram of the systematic review.

The selected studies contained 400 samples from 252 patients (97 control males, 90 schizophrenic males, 34 control females, and 31 schizophrenic females) (**Supplementary Table 2**). We performed an exploratory analysis that revealed the absence of biases or batch effects in the samples from all studies except for GSE42546, in which we found one outlier in the schizophrenic male group.

### Meta-analysis and Functional Enrichment

#### 1. Prefrontal Cortex

We meta-analyzed 27466 genes from the PFC, revealing the significantly altered expression of 265 genes in schizophrenia patients (44 increased in females and 221 increased in males) (**Figure 2A**). From these 265 genes, we found 90 genes (34%) with previously reported associations with schizophrenia according to the Open Targets database (**Supplementary Table 3**). Among them, the most affected genes upregulated in females were related to cellular signaling (GNA15, NPBWR2) and mitochondrial pseudogenes (MTND4P12, MTRNR2L8, MTCO1P12), whereas in males were related to inflammatory and immune response (CASP1, CASP8, C3, F13A1, HEPACAM) (Figure 2B).

**Figure 2.**
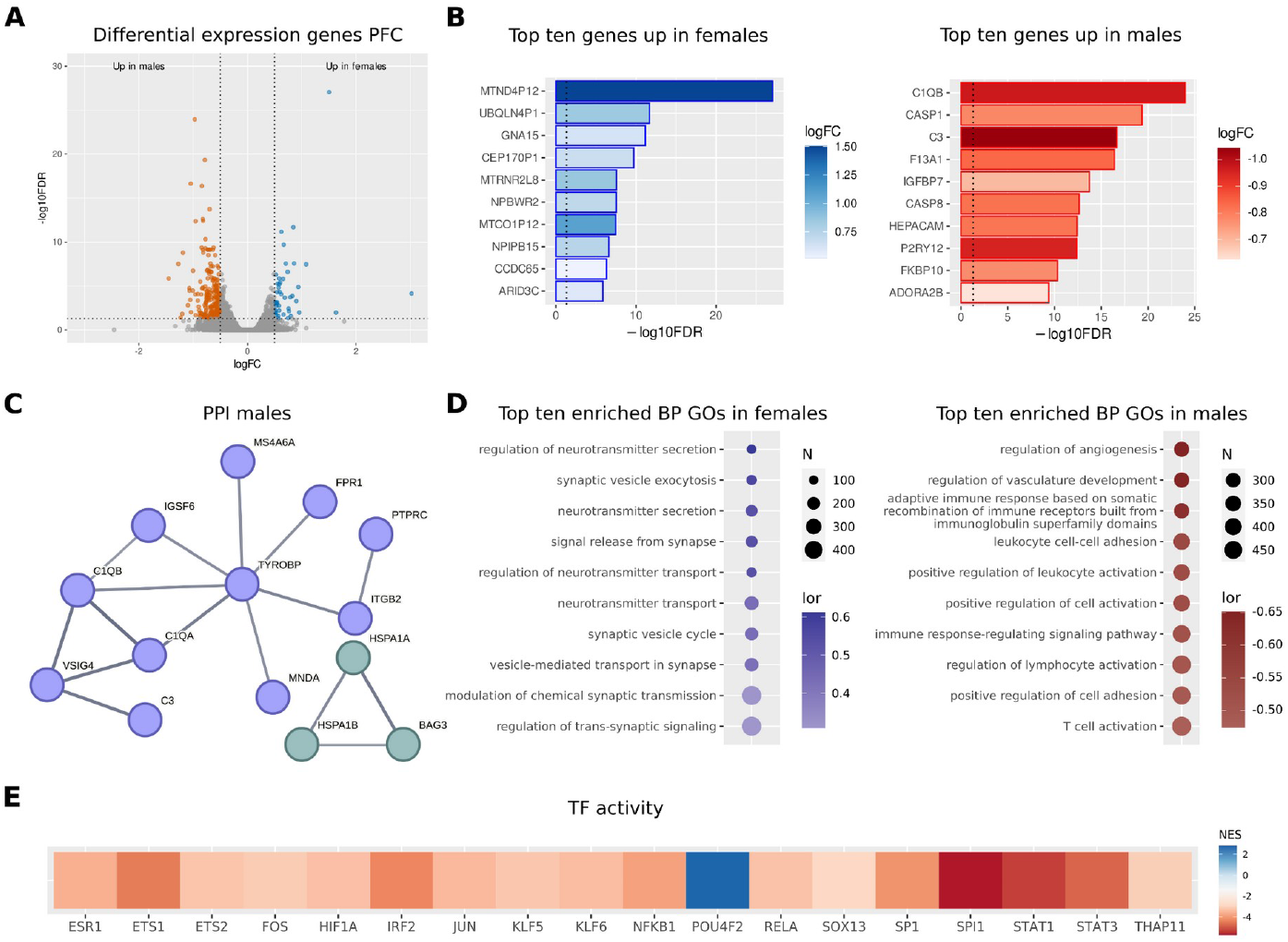
Gene expression meta-analysis in the PFC of male and female schizophrenia patients. **A**) Volcano plot depicting analyzed genes. Vertical dashed lines set at logFC 0.5 and -0.5, and horizontal dashed lines set at 1.301 -log10FDR, equivalent to a p-value of 0.05. Red dots = genes overexpressed in males, and blue dots = genes overexpressed in females. **B**) Bar plots of the top ten overexpressed genes in males (red) and females (blue). Vertical dashed lines set at p-value = 0.05. **C**) PPI network generated using genes overexpressed in males. **D**) Dot plots reporting the top ten enriched functions in males (red) and females (blue) and males (red). **E**) Heatmap showing transcription factors whose activity becomes significantly increased in males (red tones) and females (blue tones). Activation values are measured as normalized enrichment scores (NES).

Using the STRING web tool, we explored PPI networks for schizophrenia-associated DEGs in the PFC. In male schizophrenia patients, we encountered a significantly enriched network (p-value < 1.0^-16^) with two major clusters (**Figure 2C**): a central cluster (light blue, eleven nodes) related to immune response and inflammation, and another cluster (light green, three nodes) related to protein processing. In addition, we found multiple independent pairs of interacting proteins (**Supplementary Table 4**). We failed to encounter a significantly enriched network when analyzing DEGs in the PFC in female schizophrenia patients.

We next performed a GSEA on BP GO terms to assess the functional role of schizophrenia-associated genes. We found 1149 significantly affected functions in schizophrenia patients (**Figure 2D**): 57 significantly increased in females (most of them related to synapse regulation), and 1085 BP increased in males (related to immune system stimulation, extracellular matrix, and cell adhesion) (**Supplementary Table 5**).

Finally, we analyzed alterations to transcription factor activity in the PFC in schizophrenia patients (**Supplementary Table 6**) -we found an increase in the activity of seventeen transcription factors in males (ESR1, ETS1, ETS2, FOS, HIF1A, IRF2, JUN, KLF5, KLF6, NFKB1, RELA, SOX13, SP1, SPI1, STAT1, STAT3, and THAP11) and one transcription factor in females (POU4F2).

#### 2. Hippocampus

We compared 15365 genes from the hippocampus (**Figure 3A**), revealing 1414 significant DEGs in schizophrenic patients (765 increased in females and 649 increased in males). From these 1414 genes, we found 478 genes (34%) with previously reported associations with schizophrenia, according to the Open Targets database (**Supplementary Table 7**). Among them, the most affected genes upregulated in females were related to inflammation and immune system activation (CERCAM, CSF1R, IL1A, IL1B, ITGAX, TNFAIP3), whereas in males were related to multiple functions (Figure 3B).

**Figure 3.**
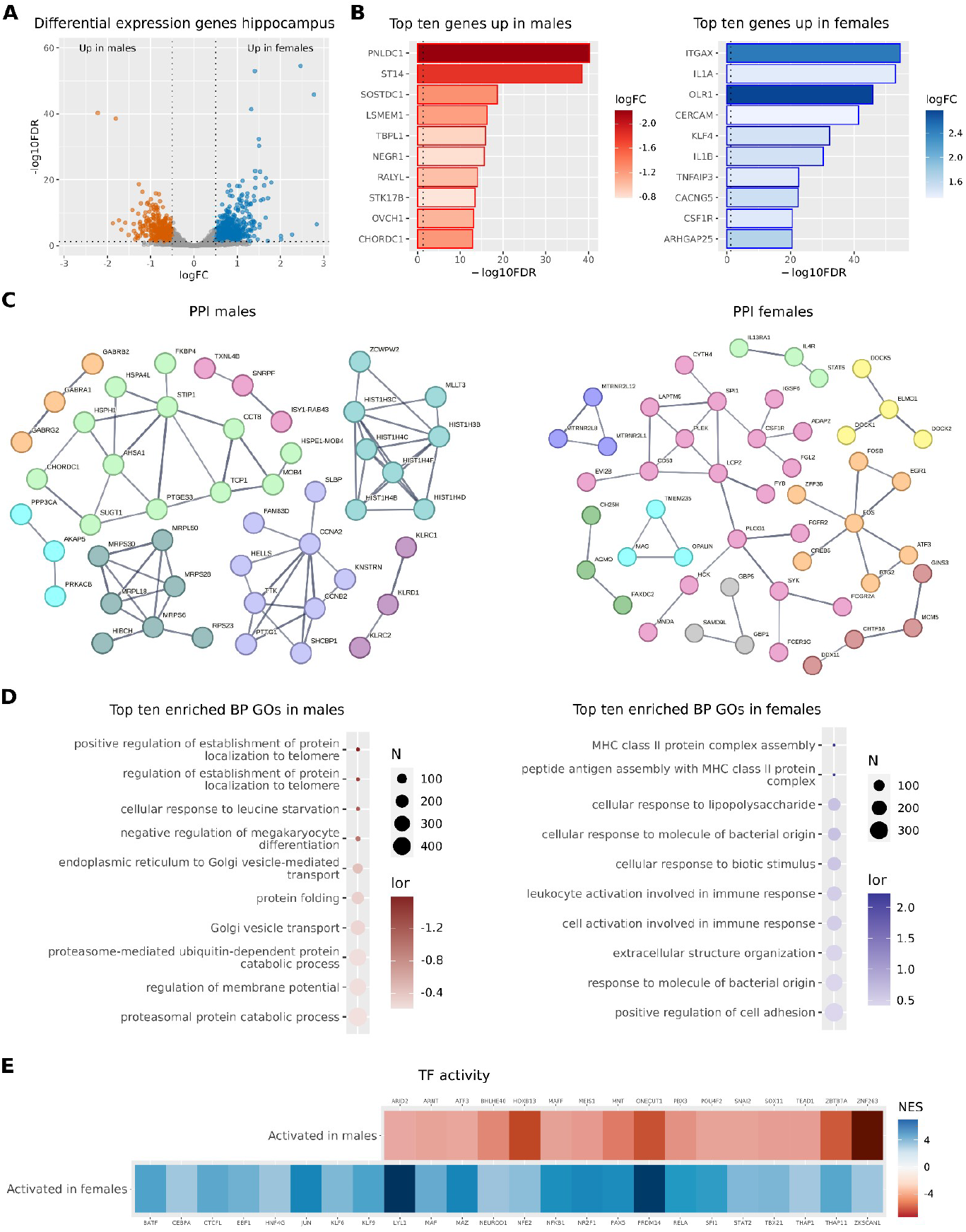
Gene expression meta-analysis in the hippocampus of male and female schizophrenia patients. **A**) Volcano plot depicting the genes analyzed. Vertical dashed lines set at logFC 0.5 and -0.5, and horizontal dashed lines set at 1.301 -log10FDR, equivalent to a p-value of 0.05. Red dots = genes overexpressed in males, and blue dots = genes overexpressed in females. **B**) Bar plots of the top ten overexpressed genes in males (red) and females (blue). Vertical dashed lines set at p-value = 0.05. **C**) PPI network generated using genes overexpressed in males (left panel) and females (right panel). **D**) Dot plots reporting the top ten enriched functions in males (red) and females (blue) and males (red). **E**) Heatmap showing transcription factors whose activity becomes significantly increased in males (red tones) and females (blue tones).

Analysis of PPI networks for DEGs in the hippocampus revealed significantly enriched networks in male (p-value < 1.0^-16^) and female (p-value < 2.3^-5^) schizophrenia patients (**Figure 3C**). In males, the leading cluster (pink, nineteen nodes) related to immune response; other significant clusters related to cytoskeletal rearrangement associated with immune response (yellow, four nodes), transcription activation (orange, seven nodes), oligodendrocyte activity (light blue, three nodes), and interleukin activation (light green, three nodes and grey, three nodes). In females, we found four principal clusters related to chaperone components (light green, twelve nodes), multiple mitochondrial ribosomal proteins (dark green, seven nodes), histone cluster 1 components (light blue, eight nodes), and proliferation-related proteins (light purple, nine nodes). In the hippocampus of male (**Supplementary Table 8**) and female (**Supplementary Table 9**) schizophrenia patients, we also encountered multiple pairs of interacting proteins.

We performed a GSEA on these genes and found 238 significantly affected functions in schizophrenia patients (218 increased in females and 20 increased in males) (**Figure 3D, Supplementary Table 10**). Interestingly, we observed increased functions related to immune system activation, cell adhesion, and extracellular matrix in females (opposite to that found in the PFC), suggesting that schizophrenia impacts the PFC and the hippocampus differently according to sex. The functions increased in males primarily related to the processing, locating, and regulating of protein synthesis and folding.

Finally, analysis of altered transcription factor activity in the hippocampus in schizophrenia patients revealed the increased activity of sixteen transcription factors in males (ARID2, ARNT, ATF3, BHLHE40, HOXB13, NAFF, MEIS1, MNT, ONECUT1, PBX3, POUF2, SNAI2, SOX11, TEAD1, ZBTB7A, and ZNF263) and twenty-four transcription factors in females (BATF, CEBPA, CTCFL, EBF1, HNF4G, JUN, KLF6, KLF9, LYL1, MAF, MAZ, NEUROD1, NFE2, NFKB1, NR2F1, PAX5, PRDM14, RELA, SPI1, STAT2, TBX21, THAP1, THAP11, and ZKSCAN1) (**Supplementary Table 11**).

#### 3. Whole Brain Analysis

We finally performed a global meta-analysis of 29005 genes that included all data sets, reflecting the whole brain. We encountered 66 significant DEGs in schizophrenic patients (25 increased in females and 41 increased in males) (**Figure 4A**). From these 66 genes, we found 17 (26%) with previously reported associations with schizophrenia, according to the Open Targets database (**Supplementary Table 12**). Among them, the most affected genes upregulated in females were long non-coding genes (LOC101928251, LOC101929719) and pseudogenes (MTND1P23, UBQLN4P1, CEP170P1), whereas in males related to multiple functions (**Figure 4B**). Analysis of PPI networks for schizophrenia-associated DEGs in the whole brain failed to reveal any significantly enriched PPI networks for male or female patients.

**Figure 4.**
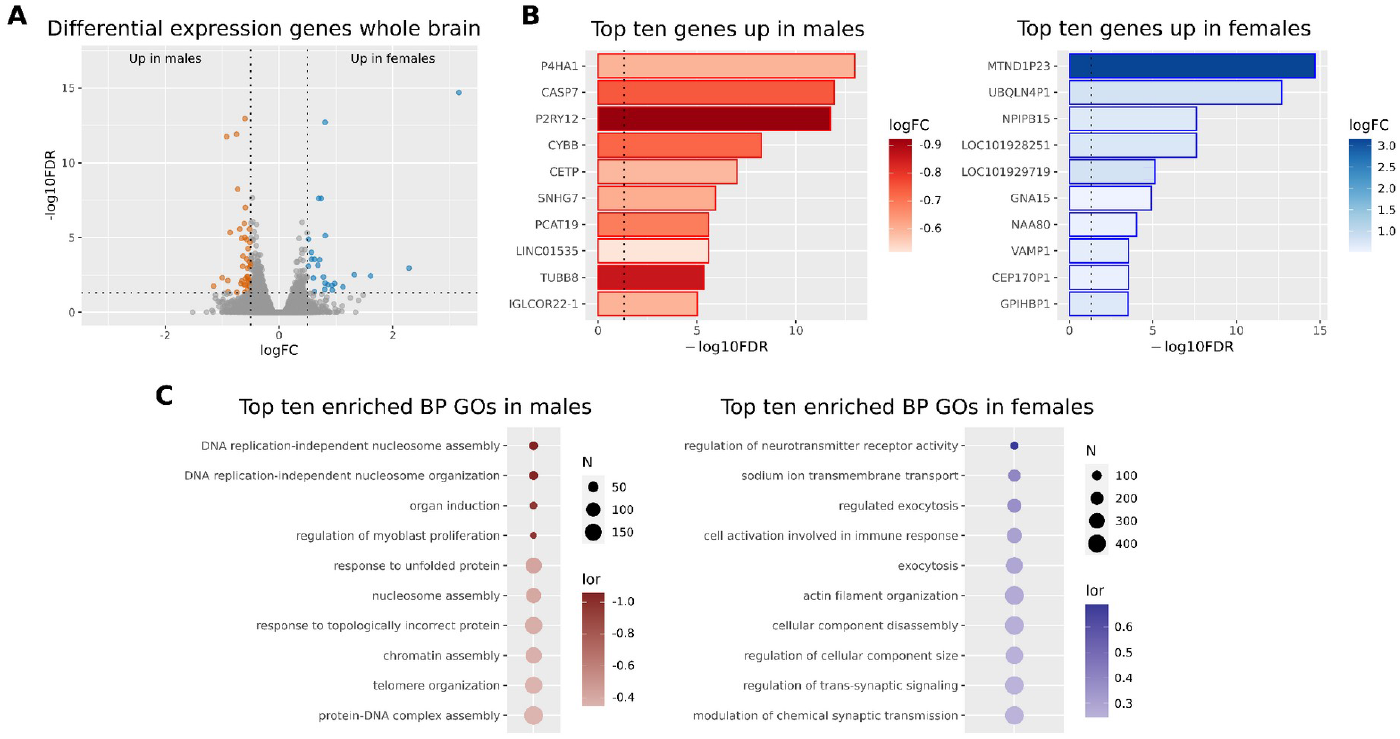
Gene expression meta-analysis in the whole brain of male and female schizophrenia patients. **A**) Volcano plot depicting the genes analyzed. Vertical dashed lines set at logFC 0.5 and -0.5, and horizontal dashed lines set at 1.301 -log10FDR, equivalent to a p-value of 0.05. Red dots = genes overexpressed in males, and blue dots = genes overexpressed in females. **B**) Bar plots of the top ten overexpressed genes in males (red) and females (blue). Vertical dashed lines set at p-value = 0.05. **C**) PPI network generated using genes overexpressed in males (left panel) and females (right panel).

GSEA on these genes in the whole brain revealed 44 significantly affected functions (34 increased in females and ten increased in males) (**Figure 4C, Supplementary Table 13**). Similarly to the PFC, we observed increased functions related to synaptic transmission in female and functions related to protein folding and nucleosome organization in male schizophrenic patients.

### The intersection of PFC, Hippocampus, and Whole Brain Meta-analyses

We finally explored the intersection of the significant DEGs and altered functions in the meta-analyses performed in schizophrenic male and female patients (**Figure 5**). We encountered most intersections between meta-analyses for genes overexpressed in the same group (i.e., genes increased in males in the hippocampus and PFC); however, we found the most significant intersection between genes overexpressed in the female hippocampus and male PFC in schizophrenic patients (*VEGCF, FYCO1, GIMAP8, SLC2A5, ARHGDIB, CMTM7, CPQ, IGS6, ELK3, RNF122, TNFRSF1B, ARHGAP29, HEY2, ACSBG1*, and *MNDA*) (**Figure 5A**). This finding highlights common schizophrenia-associated alterations in the transcriptomic profile of distinct regions. We explored the potential interactions of these genes using PPI enrichment; however, we did not find significantly enriched networks. Analysis of functional enrichment again revealed an overlap of functions increased in the female hippocampus and male PFC in schizophrenic patients (**Figure 5B, Supplementary Table 14**), supporting the idea of common schizophrenia-associated alterations in these two distinct regions.

**Figure 5.**
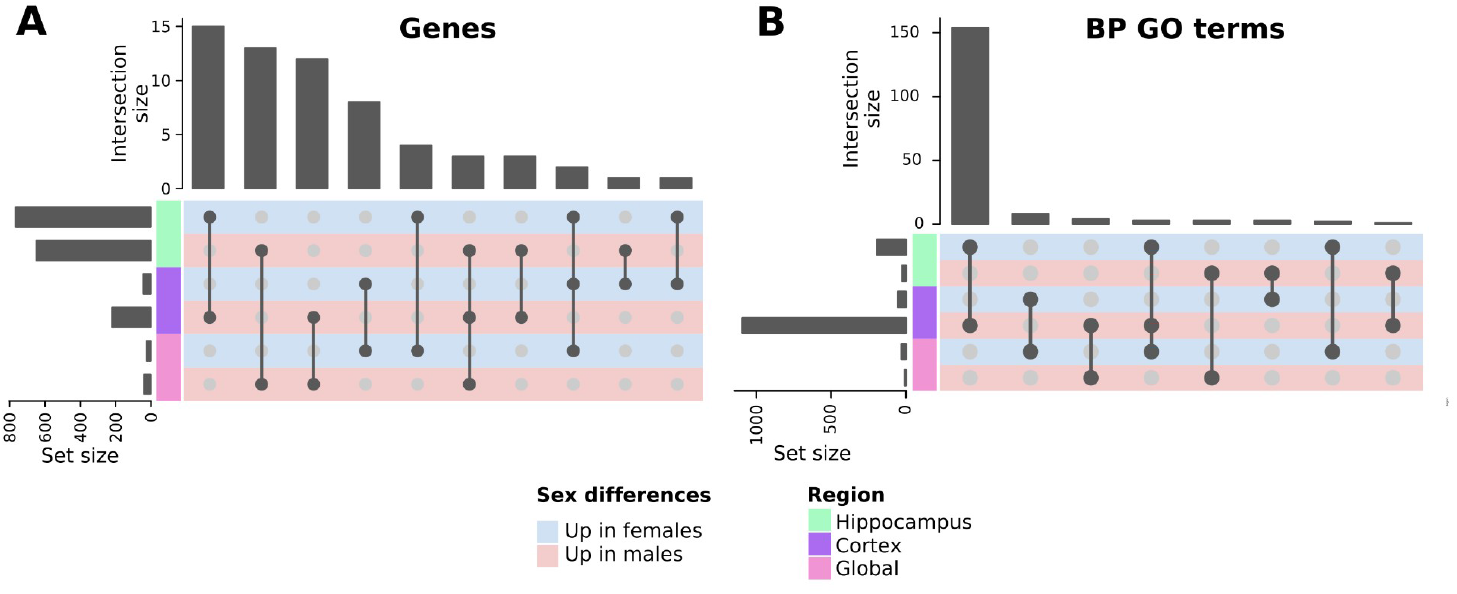
The intersection of PFC, Hippocampus, and Whole Brain Gene Expression Meta-analyses, and BP GO terms in Male and Female Schizophrenic Patients. **A**) Upset plot showing the intersections of DEGs between comparisons (light blue rows = genes overexpressed in females, light red rows = genes overexpressed in males) and regions (left column: light green = hippocampus meta-analysis, purple = PFC meta-analysis and pink = whole brain meta-analysis). **B**) Upset plot demonstrating the intersection of commonly dysregulated BP GO terms between comparisons (light blue rows = genes overexpressed in females, light red rows = genes overexpressed in males) and regions (left column: light green = hippocampus meta-analysis, purple = PFC meta-analysis and pink = whole brain meta-analysis).

## Discussion

Despite multiple sex-based differences described in the onset and symptomatology of schizophrenia, the molecular mechanisms that give rise to said differences remain mostly unexplored. To fill the existing knowledge gap, we reviewed RNA-seq-based gene expression studies in brain samples of schizophrenia patients and performed three meta-analyses on the PFC, hippocampus, and whole brain using a sex-based approach.

We first determined the effect of sex on gene expression in the PFC of schizophrenia patients, finding the most DEGs in males (with a third of these DEGS previously related to schizophrenia). This finding contrasts with a recent study that analyzed RNA-seq data in the DLPFC from two cohorts in which Hoffman et al. found that only the ALKBH3 gene displayed any degree of sex-based differential expression [13]; of note, this gene displayed no significant alteration in our meta-analyses. A recent systematic review focused on schizophrenia by Merikangas et al. discovered 160 DEGs when combining multiple technologies (microarray and RNA-seq) and tissues (blood, brain, and induced stem pluripotent cells, among others) from male and female patients [35]. We found ten of those DEGs also differentially expressed in our PFC meta-analysis (CASP1, GBP2, HSPA1A, IFITM2, IFITM3, LRRC37A2, NAGA, PDGFRB, RERGL, and ZC3HAV1); all displayed overexpression in males compared to females and mainly related to immune function, cortical immune activation, and inflammation in schizophrenia [35,36]. We also found the increased expression of component 4 (C4B) and C1QA and C1QB genes in male schizophrenia patients, which participate in inflammation and have been previously linked to schizophrenia vulnerability [37,38]; meanwhile, increased C4 expression also prompts microglia-mediated engulfment in mice [39]. Most top BP GO terms differentially increased in male schizophrenia patients related to immune system activation and neuroinflammation. Transcription factor activity analysis also revealed a panel of seventeen transcription factors with more significant activity in males; NFKB1 [40], FOS [41], and KLF6 [42] had been linked to schizophrenia previously, although none had been specifically linked to male schizophrenic patients. Interestingly, the top enriched BP GO terms overexpressed in female schizophrenia patients were related to synaptic function. A recent meta-analysis focused on synaptic protein changes in schizophrenia (but not sex) revealed the decreased expression of synaptophysin (SYP) in the DLPFC and cingulate cortex [43]. Here, we report the increased expression of SYP and another gene related to synaptic function (SV2C) in female schizophrenia patients. Of note, Onwordi et al. described the reduced expression of SV2A, a member of the same protein family as SV2C, in patients using positron emission tomography (PET) imaging [44]. Altogether, our results provide evidence for a scenario of significant inflammatory impairment and relative synaptic loss in male compared to female schizophrenia patients in the PFC, a previously undescribed link.

We found the most extensive set of DEGs (649 increased in males and 765 increased in females) when considering the meta-analysis carried out in the hippocampus. Comparing these results with a previously described meta-analysis [35], we found 16 common genes, three increased in male (FAM8A1, SNCA, and SPR) and 13 increased in female schizophrenia patients (ANXA4, CD82, CDKN1A, GNG7, HIPK2, ITGA5, NR4A1, NUAK2, PILRB, PPP1R15A, RRBP1, SRGN, and TBX2). Most of this latter set of genes play roles related to the immune system. Additionally, GSEA results demonstrated an increased number of BP terms related to immune responses in female schizophrenia patients in the hippocampus. Several studies have linked hippocampal neuroinflammation with schizophrenia [35,36], although any sex bias remains unexplored. We also found seven common transcription factors that displayed more significant activity in the PFC of male and the hippocampus of female schizophrenia patients (JUN, KLF6, NFKB1, POU4F2, RELA, SPI1, and THAP11), suggesting the common effects of increased activation of the immune system and neuroinflammation in both regions. Meanwhile, our results reported that the top BP terms displaying an increase in the hippocampus of male schizophrenia patients relate to protein folding and the ubiquitin-proteasome system [45]. Nucifora et al. previously reported an increase in protein insolubility in schizophrenia [46], while Nishimura et al. described an increase in ubiquitin immunoreactivity in the hippocampus in schizophrenia patients (primarily males) [47]. Interestingly, we found the increased expression of four ubiquitin-conjugating enzymes (E2) -UBED1, UBED3, UBEB, and UBE2U -in male schizophrenia patients. Studies have previously linked altered gene expression related to E2 activity in the hippocampus to schizophrenia (without a sex perspective) [48,49]. Therefore, our results support previous findings regarding schizophrenia-associated effects in the hippocampus and identify differentiated outcomes based on patient sex.

Our subsequent global meta-analysis included two additional studies of the amygdala and nucleus accumbens in schizophrenia patients; we encountered 41 DEGs with higher expression in males and 25 in females. The heat shock protein 70 (HSPA1A, increased in males) overlaps with the findings of a previous systematic review [35], highlighting the impairment of protein processing in schizophrenia. From total 66 DEGs, we identified 11 (16%) as pseudogenes, which comprised two main groups: mitochondrial pseudogenes increased in females (MTND1P23, MTCO1P12, MTCO1P40, and MTND2P28) and ribosomal pseudogenes increased in male schizophrenia patients (RPS4XP22, RPS16P4, and RPS28P7). Although the role of pseudogenes in schizophrenia remains relatively unexplored [50], these two clusters suggest a characteristic sex-based effect. Additionally, 15 (23%) were non-coding RNA (ncRNA), with twelve increased in male schizophrenia patients and none previously described in other systematic reviews of schizophrenia [51]. This finding agrees with a study by Hu et al. that linked increased ncRNA expression with inflammatory processes and immune system activation in schizophrenia [33].

### Strengths and Limitations

The systematic review and meta-analysis strategy integrates multiple datasets, thereby increasing statistical power and allowing the discovery of subtle effects compared to individual studies. A range of previous studies have integrated data from numerous sources to characterize sex-based differences in the human transcriptome [52–56]; however, the analysis of sex-based differences in bulk RNA-seq data in schizophrenia have been limited to the DLPFC [13]. Our systematic review allowed us to include additional studies carried out on the hippocampus, nucleus accumbens, and amygdala, which represent regions affected by schizophrenia [57,58]. Our study does present some limitations, including bias in the sex of subjects (significantly more males than females), which constrains the statistical power of analyses. Furthermore, bulk RNA-seq studies do not allow the assignment of identified differences to specific cell populations, limiting a deeper understanding of their consequences. Finally, important covariates such as medication usage, smoking status, years of disease after diagnosis, and post-mortem interval were not included in the metadata of most studies, thus increasing the unexplained variability of the data. The addition of further data, especially from female donors and the expansion to single-cell RNA-seq technology, could improve our ability to detect cell type-specific sex-based gene expression differences associated with schizophrenia.

### Perspectives and significance

The results described support a different sex-based impact of schizophrenia in multiple brain regions. In the PFC, our data suggest a larger inflammatory impairment in males and a reduced impact in neurotransmission of females. On the other hand, the hippocampus of females seems more affected by immune system activation, whereas the impact of schizophrenia in males affects protein processing. Together, our results outline common alterations in the hippocampus of females and the PFC of males. The present study takes a novel approach to assess the sex differences described in schizophrenia through a comprehensive bioinformatic strategy.

## Conclusions

To the best of our knowledge, we present the first systematic review and meta-analysis of transcriptomic studies in schizophrenia patients using a sex-based approach. Our results describe the differential schizophrenia-associated effects in the transcriptomic profiles of male and female patients based on the brain region analyzed. These data highlight the overlapping immune activation processes in the PFC of male and the hippocampus of female schizophrenia patients. Moreover, our global meta-analysis provided evidence of the differential expression of multiple non-protein coding genes in male and female schizophrenia patients, suggesting their potential role in disease development, monitoring, and treatment.

## Acknowledgments

The authors thank the Principe Felipe Research Center (CIPF) for providing access to the cluster, co-funded by European Regional Development Funds (FEDER) in Valencian Community 2014-2020. The authors also thank Stuart P. Atkinson for reviewing the manuscript.

## Author contributions

HC and FGG conceived the project and designed the work. HC acquired, analyzed and interpreted data, and wrote the manuscript. MRH, MJE, JN, MIV, and FGG revised critically the work for and approved the final version to be published.

## Data availability

The data used for the analyses described in this work are publicly available at GEO and SRA. The accession numbers of the GEO datasets downloaded are GSE174407, GSE138082, GSE80655, GSE42546, and GSE78936. The accession number of the SRA dataset downloaded is SRP102186.

## Code availability

The code developed for the analyses described in this work has been made publicly available in Zenodo (http://doi.org/10.5281/zenodo.7778277).

## Ethics approval and consent to participate

Not applicable

## Consent for publication

Not applicable

